# Chromatin landscape dynamics in development of the plant parasitic nematode *Meloidogyne incognita*

**DOI:** 10.1101/2021.05.11.443567

**Authors:** Rahim Hassanaly-Goulamhoussen, Ronaldo De Carvalho Augusto, Nathalie Marteu-Garello, Arthur Péré, Bruno Favery, Martine Da Rocha, Etienne G.J. Danchin, Pierre Abad, Christoph Grunau, Laetitia Perfus-Barbeoch

**Affiliations:** Université Côte d’Azur, INRAE, CNRS, Institut Sophia Agrobiotech UMR 1355, France; IHPE, Univ Montpellier, CNRS, IFREMER, Univ Perpignan Via Domitia, Perpignan, France; Laboratory of Biology and Modeling of the Cell, Ecole Normale Supérieure de Lyon, CNRS, Université Claude Bernard de Lyon, Université de Lyon, Lyon, France

**Keywords:** Histone modifications, Epigenetics, Root-knot nematode, Development, Parasitism.

## Abstract

In model organisms, epigenome dynamics underlies a plethora of biological processes. The role of epigenetic modifications in development and parasitism in nematode pests remains unknown. The root-knot nematode *Meloidogyne incognita* adapts rapidly to unfavorable conditions, despite its asexual reproduction. However, the mechanisms underlying this remarkable plasticity and their potential impact on gene expression remain unknown. This study provides the first insight into contribution of epigenetic mechanisms to this plasticity, by studying histone modifications in *M. incognita*. The distribution of five histone modifications revealed the existence of strong epigenetic signatures, similar to those found in the model nematode *Caenorhabditis elegans*. We investigated their impact on chromatin structure and their distribution relative to transposable elements (TE) loci. We assessed the influence of the chromatin landscape on gene expression at two developmental stages: eggs, and pre-parasitic juveniles. H3K4me3 histone modification was strongly correlated with high levels of expression for protein-coding genes implicated in stage-specific processes during *M. incognita* development. We provided new insights in the dynamic regulation of parasitism genes kept under histone modifications silencing. In this pioneering study, we establish a comprehensive framework for the importance of epigenetic mechanisms in the regulation of the genome expression and its stability in plant-parasitic nematodes.

**Author summary:** The nematode *Meloidogyne incognita* is one of the most destructive plant parasites worldwide. Its ability to infect a wide range of hosts and its high adaptability contribute to its parasitic success. We investigated the role of epigenetic mechanisms — specifically post-translational histone modifications — in the parasitic life cycle. We showed these modifications are linked to gene expression regulation and likely contribute to nematode development and pathogenicity.

## Introduction

Crops are continually attacked by a wide range of pests and parasites. Plant-parasitic nematodes are thought to be one of the main causes of damages in food crops, resulting in yield losses of more than $150 billion worldwide [1]. Root knot nematodes (RKN), *Meloidogyne spp*, are among the most rapidly spreading of all crop pests and pathogens [2]. Their rapid spread may have been facilitated by their wide host range, high fecundity, and parthenogenetic reproduction, allowing infestations to become established with relatively few individuals [1]. Understanding the determinants of the extreme adaptive capacity of RKN is crucial for the development of effective and sustainable control methods.

*Meloidogyne incognita* is the most ubiquitous RKN with an obligate biotroph lifestyle. It feeds exclusively on living cells within the vascular cylinder of the root [3]. The freshly hatched second-stage pre-parasitic juveniles (J2s) within the soil are attracted to the root tip of the host plant. These microscopic J2s (400 μm long and 15 μm wide) invade host roots close to the root elongation zone, through the physical and enzymatic destruction of plant cell walls in the root epidermis, eventually reaching the vascular cylinder, where they establish a permanent feeding site [4]. To this end, infective juveniles secrete molecules known as effectors, to induce major cellular changes in recipient host cells and evade plant defense responses. These effector proteins are translocated directly from the secretory gland cells into the host cells by a syringe-like structure, called stylet [5]. The tissue around the permanent feeding site typically shows signs of hyperplasia, resulting in the characteristic knot-like shape of roots infected with RKN. Once they begin feeding, the J2s become sedentary, going through three molts before becoming mature adults. The females release eggs onto the root surface, and embryogenesis within the eggs is followed by the first molt, generating second-stage juveniles. Males are produced in unfavorable conditions (e.g., resistant host), and they migrate out of the plant without developing further and without playing a role in reproduction [6].

Despite its mitotic parthenogenetic mode of reproduction, presumably resulting in low genetic plasticity, *M. incognita* can adapt rapidly to unfavorable conditions [7, 8]. The mechanisms underlying this adaptability have yet to be elucidated. Population genomics analyses have revealed only low genome variability at the SNP level between *M. incognita* isolates across the globe [8]. Furthermore, these point mutations did not correlate with the ranges of compatible plant host species. A follow-up population genomics study on Japanese isolates [9] confirmed the low genome variability at the SNP level but identified some correlations with infection compatibility of different cultivars of the same plant species (sweet potato). Taken together, these studies suggest point mutations are not the sole genome plasticity factors involved in the adaptive evolution of *M. incognita*. Consequently, other genome plasticity factors have also been investigated in this species, including movements of transposable elements (TE) and gene copy number variations (CNV). High similarity between TE copies and their consensus sequences suggest they have been recently active in the *M. incognita* genome [10]. Studying variations of their frequencies across geographical isolates allowed identification of isolate-specific TE insertions, including in coding or regulatory regions, suggesting TE movements might constitute a genome plasticity factor with functional consequences. However, no evidence yet for an adaptive role of these movements were shown in this species and nothing is known about the mechanisms underlying their regulation or amplification. In addition, convergent gene CNV have been shown to correlate with rapid breaking down of tomato plant resistance, suggesting an adaptive role, although causative relation has not yet been shown [11] and the underlying mechanisms are also unknown. Because a strategy to explain *M. incognita*’s capacity to adapt in a fast-fluctuating environment is lacking, investigating whether epigenetic mechanisms do occur and have possible impact on genome regulation is timely. Indeed, the epigenetic control of transposable elements has been identified as an important factor of genome evolution [12]. Furthermore, the epigenome dynamics of multicellular organisms are associated with transitions in cell cycle development, germline specification, gametogenesis, and inheritance. Within the cell, nuclear DNA is packaged and ordered into chromatin by histone proteins [13, 14]. Chromatin can adopt different conformational states directly influencing gene expression, from relaxed transcriptionally active euchromatin to condensed transcriptionally inactive heterochromatin. Specific enzymes regulate histone structure and function through chemical modifications to the histone proteins, such as acetylation and methylation. In many organisms, euchromatin displays an enrichment in the di-(or tri-) methylation of the lysine 4 residue of histone 3 (H3K4me3), whereas heterochromatin displays enrichment in the trimethylation of the lysine 9 or lysine 27 residue of histone 3 (H3K9me3 and H3K27me3) [15]. Specific combinations of histone modifications are associated with transcriptionally permissive or repressive chromatin structures, thereby controlling gene expression at the transcriptional level [16]. Other organisms, such as *Saccharomyces cerevisiae*, display an unusual regulation of histone modifications, with a lack of H3K9me3 modification and the establishment of alternative modifications defining the silent state of chromatin [17].

Chromatin immunoprecipitation followed by high-throughput sequencing (ChIP-seq) is a powerful method for generating genome-wide maps of interactions between proteins and DNA, including posttranslational histone modifications, and for mapping histone variants [18]. Extensive epigenetic studies have been performed in the model nematode *Caenorhabditis elegans*, addressing its functional genomic elements, including histone modifications in response to the environment [19]. Previous studies have shown that *M. incognita* lacks 5-methylcytosine (5mC) and has no cytosine-DNA (cytosine-5)-methyltransferase 1 (DNMT1) or DNMT3 [20, 21] which is similar to what is known for *C. elegans* [22]. Low-level DNA N(6)-methylation (6mA-DNA) has been identified as an alternative carrier of epigenetic information in *C. elegans* [23]. However, the physiological relevance of 6-mA-DNA remains unclear. Apart from this model species, the role of chromatin modifications has not been studied in nematodes. The studies performed to date have been limited to bioinformatics analyses indicating that potential homologs of canonical histone-modifying enzymes are conserved in the genomes of *C. elegans* and two parasitic nematodes, the food-borne animal parasite *Trichinella spiralis* and the plant parasite *M. incognita* [21, 24]. Epigenetic regulation is considered a key mechanism of parasite adaptation, and its role in plant-nematode interactions is an emerging field of study [25].

Deciphering histone modifications and their effects on gene transcription is important for understanding the key parameters underlying biological processes, including parasitic success in RKN. This study provides the first insight of the genome-wide epigenetic landscape of *M. incognita* and its direct relationship to gene transcription. Using ChIP-seq, we first analyzed the distribution of five posttranslational histone modifications. We then investigated the impact of these modifications on chromatin structure and their co-distribution relative to TE-rich regions. Finally, we assessed the influence of the chromatin landscape on gene expression during developmental, with a focus on parasitism genes, such as those encoding effectors.

## Results

### The chromatin landscape of five histone modifications in *M. incognita*

We performed ChIP-Seq analysis to study posttranslational histone modifications in *M. incognita*. We first checked the specificity of a set of commercially available antibodies and optimized the binding and sonication steps. Four out of 15 available antibodies previously used in *C. elegans* passed the two-step validation process [26]. These antibodies gave single bands on western blots and saturated signals on ChIP-titration (S1 Fig, S1 Table). They were raised specifically against H3K27ac, H4K20me1, H3K9me3 and H3K27me3, and were added to the first previously validated antibody raised against H3K4me3 [20]. ChIP-Seq data were obtained for two RKN developmental stages, eggs and pre-parasitic juveniles 2 (J2s), and were mapped to the most complete annotated *M. incognita* genome publicly available [27]. Regions displaying a specific enrichment in histone modifications were identified (S2 Fig), making study of the chromatin landscape based on these five histone modifications meaningful.

We investigated the distribution of histone modifications in the *M. incognita* genome further, by calculating the genomic frequencies of each histone modification and of the 31 histone modification combinations detected genome-wide (Table 1). These frequencies correspond to the percentage of the total genome (∼184 Mb divided by a bin size of 500 bp each) covered by each histone modification. In both eggs and J2s, H3K4me3 was the most prevalent histone modification, covering 13.9% and 14.6% of the genome, respectively. By contrast, H3K9me3, H3K27me3, H4K20me1 and H3K27ac each covered less than 4% of the genome. Very little difference in the frequencies of these modifications was observed between eggs and J2s (Table 1).

**Table 1.** Overall coverage frequencies of ChIP-Seq data.

| Histone modification combination | Whole-genome coverage in eggs (bp) | Whole-genome coverage in J2s (bp) | Proportion of whole genome in eggs (%) | Proportion of whole genome in J2s (%) | Proportion in TE in eggs (%) | Proportion in TE in J2s (%) |
| --- | --- | --- | --- | --- | --- | --- |
| [H3K4me3] | 25,235,500 | 26,575,000 | 13.861 | 14.597 | 2.899 | 3.129 |
| [H4K20me1] | 6,900,000 | 6,535,000 | 3.79 | 3.59 | 1.747 | 2.104 |
| [H3K9me3] | 6,379,000 | 6,033,500 | 3.504 | 3.314 | 8.614 | 7.108 |
| [H3K27me3] | 4,036,000 | 5,004,500 | 2.217 | 2.749 | 1.73 | 2.421 |
| [H3K9me3+H4K20me1] | 3,676,500 | 3,062,500 | 2.019 | 1.682 | 5.944 | 4.756 |
| [H3K27ac] | 2,991,500 | 2,528,000 | 1.643 | 1.389 | 0.434 | 0.386 |
| [H3K27me3+H3K9me3+H4K20me1] | 2,737,500 | 2,499,500 | 1.504 | 1.373 | 4.283 | 4.05 |
| [H3K27me3+H4K20me1] | 2,698,000 | 3,431,500 | 1.482 | 1.885 | 1.387 | 2.239 |
| [H3K27ac+H4K20me1] | 1,857,500 | 1,478,000 | 1.02 | 0.812 | 0.29 | 0.214 |
| [H3K4me3+H3K9me3] | 1,580,500 | 836,000 | 0.868 | 0.459 | 0.18 | 0.115 |
| [H3K27ac+H3K27me3+H4K20me1] | 1,519,000 | 1,643,500 | 0.834 | 0.903 | 0.362 | 0.434 |
| [H3K4me3+H4K20me1] | 1,263,500 | 1,263,000 | 0.694 | 0.694 | 0.194 | 0.168 |
| [H3K27ac+H3K4me3] | 685,000 | 624,500 | 0.376 | 0.343 | 0.091 | 0.082 |
| [H3K27me3+H3K9me3] | 657,500 | 656,000 | 0.361 | 0.36 | 0.643 | 0.617 |
| [H3K27ac+H3K27me3+H3K9me3+H4K20me1] | 545,000 | 403,500 | 0.299 | 0.222 | 0.792 | 0.379 |
| [H3K27ac+H3K27me3] | 481,000 | 544,500 | 0.264 | 0.299 | 0.096 | 0.089 |
| [H3K27ac+H3K4me3+H4K20me1] | 480,000 | 439,000 | 0.264 | 0.241 | 0.053 | 0.05 |
| [H3K4me3+H3K9me3+H4K20me1] | 258,000 | 164,500 | 0.142 | 0.09 | 0.098 | 0.06 |
| [H3K27ac+H3K9me3+H4K20me1] | 251,000 | 221,000 | 0.138 | 0.121 | 0.238 | 0.072 |
| [H3K27me3+H3K4me3] | 194,500 | 188,500 | 0.107 | 0.104 | 0.031 | 0.022 |
| [H3K27ac+H3K9me3] | 149,000 | 246,000 | 0.082 | 0.135 | 0.034 | 0.038 |
| [H3K27ac+H3K4me3+H3K9me3] | 108,500 | 67,000 | 0.06 | 0.037 | 0.019 | 0.017 |
| [H3K27ac+H3K4me3+H3K9me3+H4K20me1] | 109,500 | 76,500 | 0.06 | 0.042 | 0.043 | 0.024 |
| [H3K27ac+H3K27me3+H3K4me3+H3K9me3+H4K20me1] | 94,500 | 75,000 | 0.052 | 0.041 | 0.154 | 0.086 |
| [H3K27ac+H3K27me3+H3K4me3+H4K20me1] | 91,000 | 109,500 | 0.05 | 0.06 | 0.014 | 0.022 |
| [H3K27me3+H3K4me3+H4K20me1] | 63,000 | 76,500 | 0.035 | 0.042 | 0.007 | 0.017 |
| [H3K27me3+H3K4me3+H3K9me3+H4K20me1] | 55,000 | 28,000 | 0.03 | 0.015 | 0.036 | 0.046 |
| [H3K27ac+H3K27me3+H3K9me3] | 49,000 | 92,500 | 0.027 | 0.051 | 0.017 | 0.026 |
| [H3K27me3+H3K4me3+H3K9me3] | 44,000 | 21,500 | 0.024 | 0.012 | 0.017 | 0.007 |
| [H3K27ac+H3K27me3+H3K4me3] | 25,500 | 33,000 | 0.014 | 0.018 | 0.007 | 0.014 |
| [H3K27ac+H3K27me3+H3K4me3+H3K9me3] | 6,500 | 13,500 | 0.004 | 0.007 | 0 | 0 |
| **Total** | **65,222,000** | **64,970,500** | **35.825** | **35.687** | **30.454** | **28.792** |

Overall alignment (bp) and genomic coverage percentage of H3K4me3, H3K9me3, H3K27ac, H3K27me3 and H4K20me1 histone modifications and their combinations over the whole *M. incognita* genome and the Transposable Element annotations (TE), for Egg and J2 samples. *M. incognita* genome has been fragmented *in silico* into 500 bp bins on which histone modification enrichment was predicted with a posterior probability > 0.5. If a histone modification was predicted, the corresponding 500bp bin was counted. Coverage frequencies were calculated based on 184 Mb total *M. incognita* genome size. Three biological replicates of *M. incognita* eggs and J2s have been treated jointly to identify common histone modification enrichment using ChromstaR.

Histone modifications can act together in a combinatorial manner to exert different effects on the genome. The most frequent histone combinations observed in both eggs and J2s involved H4K20me1+H3K27me3, or H4K20me1+H3K9me3, or H4K20me1+H3K27me3+H3K9me3, with frequencies ranging between 1.3% and 2%. The other 23 combinations presented relatively low coverage, with a frequency of less than 1%. In total, ∼35% of the *M. incognita* genome was covered by the five histone modifications and their combinations (Table 1). Overall, these results reveal a consistent chromatin landscape during *M. incognita* eggs-to-J2s transition based on the five post translational histone modifications considered here.

### *M. incognita* displays canonical distribution for histone modifications

We used ChromstaR [28] to analyze the spatial pattern of statistically significant enrichment in each histone modification associated with different functional genomic elements in *M. incognita*. These associations provide clues for the functions and regulatory mechanisms of histone modifications. Spatial enrichment was calculated and represented as a heatmap for both eggs and J2s (Fig 1 and S3 Fig, respectively). Enrichment level was calculated for all the available annotations for the *M. incognita* genome: coding sequence (CDS), exon, 5’-untranslated region (UTR), messenger RNA (mRNA), non-coding RNA (ncRNA), ribosomal RNA (rRNA), TE, 3’-UTR and tRNA.

**Fig 1.**
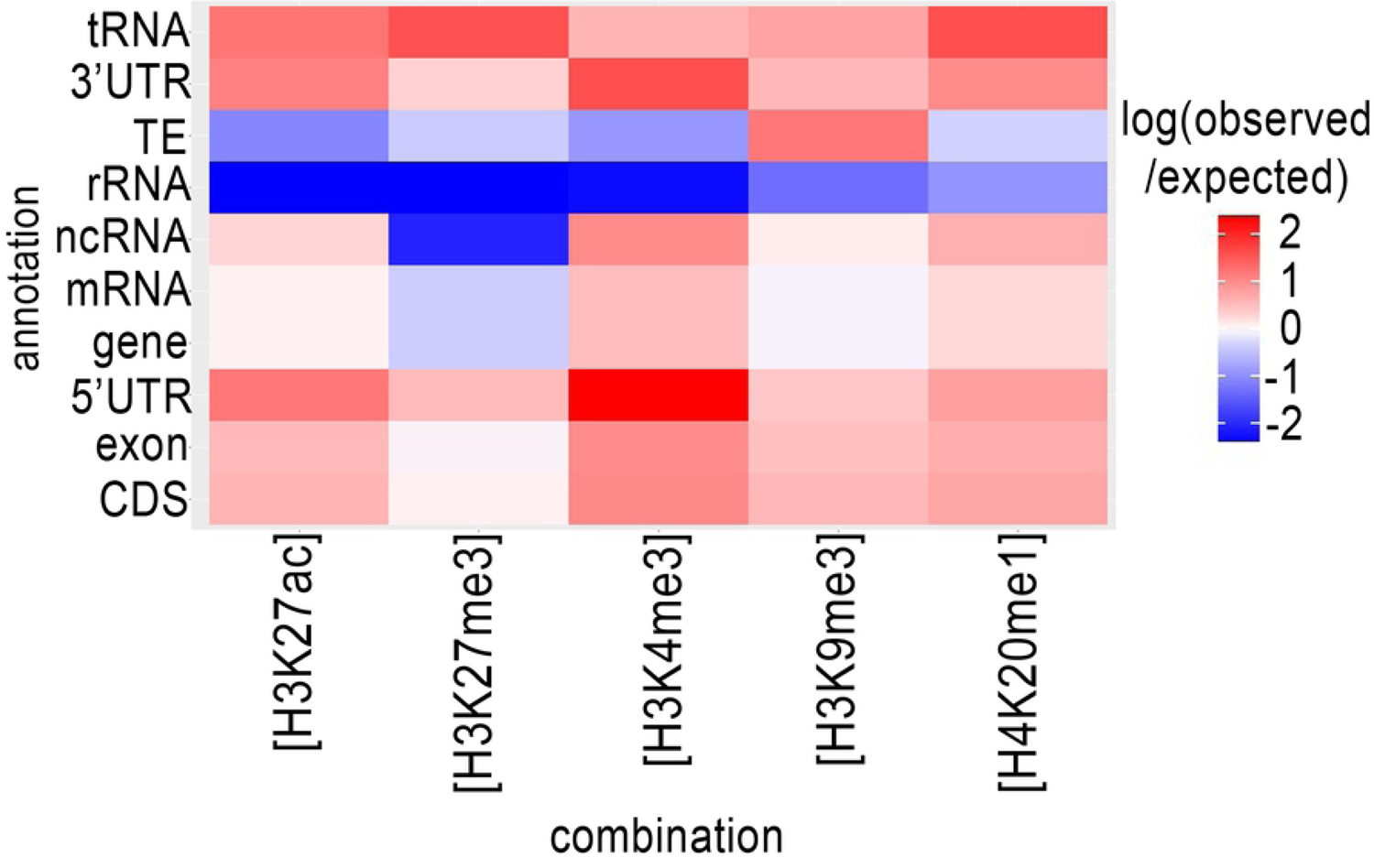
Genome wide distribution of histone modifications in relation to annotations for the *M. incognita* genome. The distribution of histone modifications was analyzed with ChromstaR, which calculated the spatial enrichment in histone modifications for different available genomic annotations. Enrichment is represented as the log(observed/expected) value and ranges from 2 (highly enriched, red) to −2 (depletion, blue). This enrichment heatmap is a 5×10 matrix representing 5 histone modifications (H3K4me3, H3K9me3, H3K27ac, H3K27me3 and H4K20me1) and 10 genomic annotated elements (CDS, exon, five prime UTR, gene, mRNA, ncRNA, rRNA, TE, three prime UTR and tRNA). Three biological replicates of *M. incognita* eggs have been treated jointly to identify common histone modification enrichment.

We observed a highly significant enrichment in H3K4me3 for sequences annotated as related to protein-coding genes (CDS, exon, UTRs and mRNA) and various types of non-protein-coding RNA genes (ncRNA and tRNA), this enrichment being strongest for the 5’-UTR. An enrichment of H3K27me3 was also observed in the 5’-UTR, however the enrichment in this modification was weak for other gene-related annotations. H3K4me3 modifications were observed less frequently than expected for rRNA and TE. H3K9me3 enrichment was observed for almost all genomic annotations, particularly for TE, but not for rRNA. Interestingly, rRNA genes displayed a relative depletion for all five histone modifications. Finally, the levels of enrichment in H3K27ac, H3K27me3 and H4K20me1 were highest for tRNA genes. A similar enrichment distribution was observed in J2s (S3 Fig).

Histone modifications associated with genomic elements were visualized on the longest scaffold, Minc3s00001, as an example (Fig 2). For H3K4me3, sharp peaks overlapping with the transcriptional start site (TSS/5’-UTR) were observed. For H3K9me3, peaks overlapping both protein-coding genes and TEs were observed, whereas H3K27ac, H3K27me3 and H4K20me1 yielded broad shapes and distributions. The distribution and enrichment patterns of histone modifications suggest a canonical role of H3K9me3 in TE repression and of H3K4me3 in promoting protein-coding gene expression (S4 Fig).

**Fig 2.**
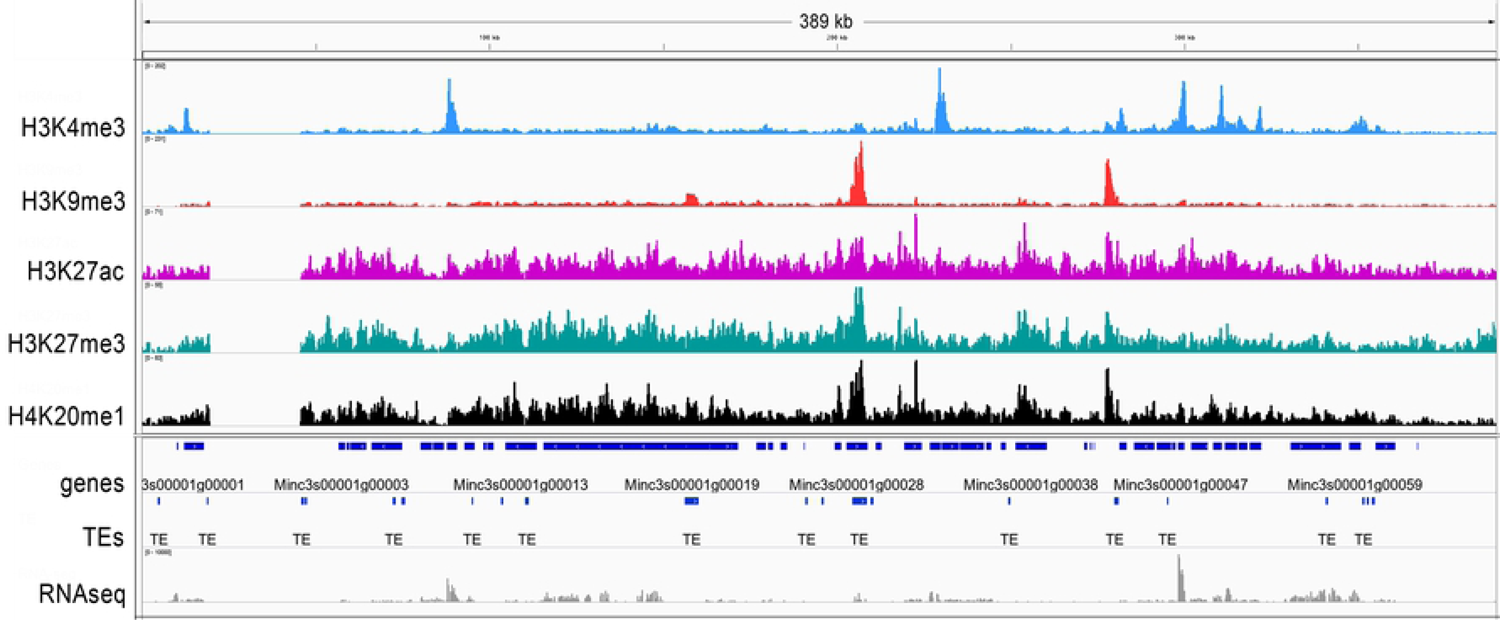
H3K4me3, H3K9me3, H3K27ac, H27me3 and H4K20me1 histone modifications on the *M. incognita* genome. The general tracks of histone modifications are illustrated on the longest scaffold (Minc3s00001) of the *M.incognita* genome. Sequence reads for H3K4me3 (blue), H3K9me3 (red), H3K27ac (pink), H3K27me3 (green) and H4K20me1 (black) samples were visualized in IGV software. Values shown on the *y* axis represent the relative enrichment of ChIP-Seq signals obtained with PeakRanger (peaks correspond to read counts after background/input subtraction). The tracks were overlaid with the *M. incognita*’s annotations (dark blue) of Genes and Transposable Elements (TE), as well as RNA-seq reads (grey). Each track contains information from one biological replicate of eggs.

### Transposable element orders display preferential enrichment in H3K9me3

TEs are important drivers of genomic plasticity in *M. incognita* [10]. The genome-wide annotation of *M. incognita* TEs identified retrotransposons and DNA-transposons, such as terminal inverted repeats (TIR), miniature inverted repeat transposable elements (MITEs), helitrons, maverick elements, long terminal repeats (LTR), long and short interspersed nuclear elements (LINE and SINE), terminal-repeat retrotransposons in miniature (TRIM), and large retrotransposon derivatives (LARD) [10]. We calculated the frequency of histone modifications associated with TE annotations (Table 1). In both eggs and J2s, H3K9me3 had the highest frequency, covering 8.6% and 7.1% of annotated TEs, respectively. By contrast, H3K4me3, H4K20me1, H3K27me3 and H3K27ac had lower frequencies, ranging from 0.3 to 2.9%. Three histone modification combinations, involving H4K20me1, were also present at a high frequency (1.4% to 5.9%) at annotated TEs. The other 23 histone combinations covered less 1% of the annotated TE. We found that H3K9me3 was observed more frequently than expected in association with all TE orders except SINE (Fig 3). H4K20me1 modification was observed more frequently in four TE orders (TRIM, MITE, TIR and helitron). By contrast, H3K4me3, H3K27ac and H3K27me3 displayed a lower association with all TE orders. The enrichment of most TE subfamilies in H3K9me3 supports the hypothesis of a role for this histone modification in repressing TE, consistent with conservation of the canonical role of this modification in *M. incognita*.

**Fig 3.**
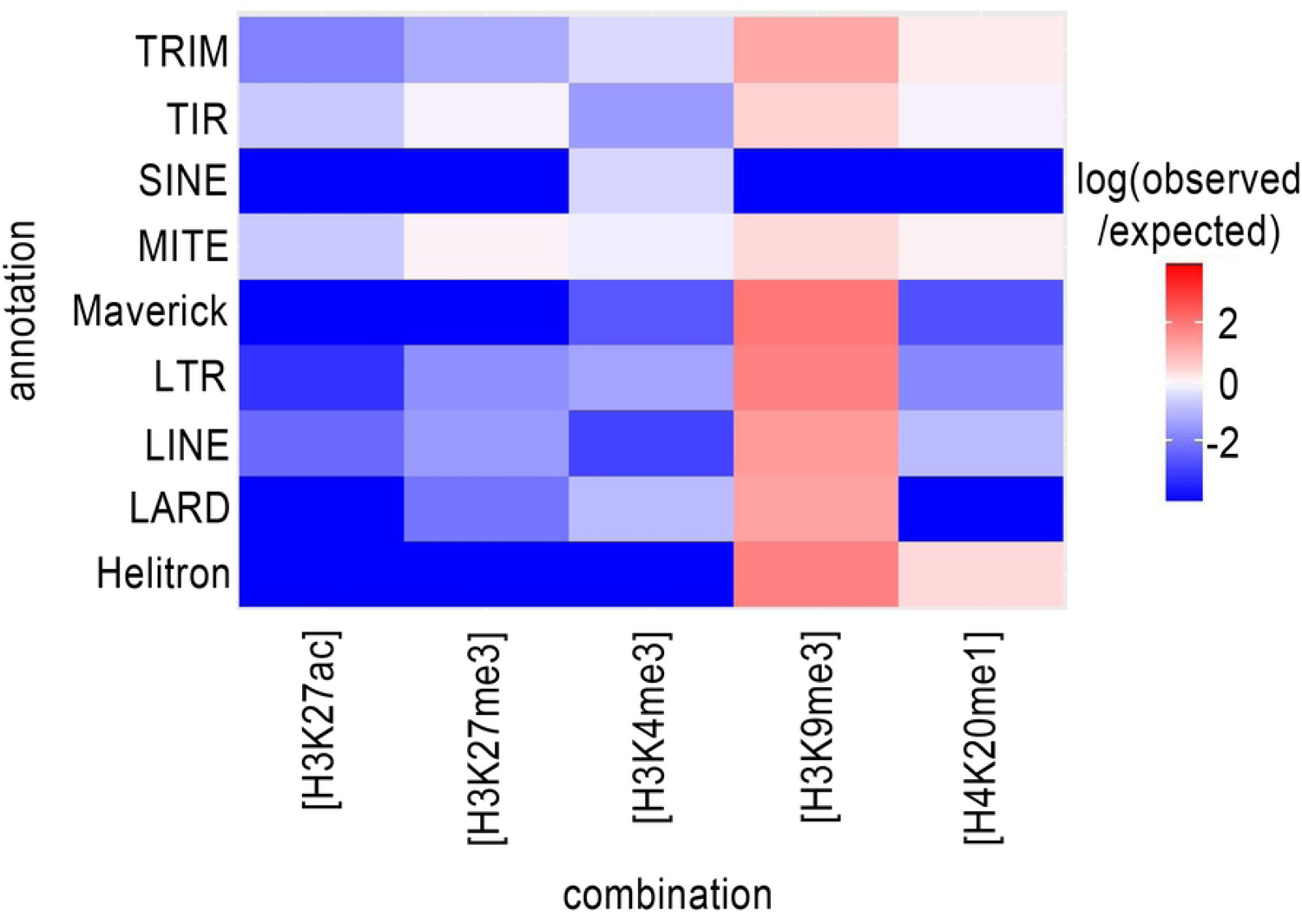
Distribution of histone modifications in relation to transposable element (TE) orders. The distribution of histone modifications was analyzed with ChromstaR, which calculated the spatial enrichment in histone modifications for annotated subfamilies of TE in *M. incognita*. Enrichment is represented as the log(observed/expected) value and ranges from 2 (highly enriched, red) to −2 (depletion, blue). This enrichment heatmap is a 5×11 matrix representing 5 histone modifications (H3K4me3, H3K9me3, H3K27ac, H3K27me3 and H4K20me1) and 11 TE orders (4 DNA-transposons (Helitron, Maverick, TIR, MITE), and 5 RNA transposons (LINE, LTR, SINE, LARD and TRIM)). Three biological replicates of *M. incognita* eggs have been treated jointly to identify common histone modification enrichment.

### The H3K4me3 modification is associated with higher levels of gene expression during nematode development

Histone modifications are known to regulate the spatiotemporal expression of protein-coding genes [29], and, thus, developmental processes. Early in the development of *M. incognita*, the transition from eggs to J2s constitutes a dramatic change in environment for the nematode, because the mobile J2s are released into the soil after hatching. We evaluated changes in both the pattern of histone modifications and gene expression during this transition, by determining the number of protein-coding genes overlapping each area of enrichment in particular histone modifications and their combinations (Table 2). We found that 13,322 of the 43,718 annotated *M. incognita* protein-coding genes were associated with at least one histone modification in eggs, whereas 23,470 genes were associated with at least one histone modification in J2s. At both developmental stages, H3K4me3 modification was associated with the largest number of genes (6,014 in eggs and 10,564 in J2s), followed by H3K9me3, H4K20me1, H3K27me3 and H3K27ac. The most prevalent histone modification combinations were the H3K9me3+H4K20me1 combination in eggs, which was associated with 531 genes, and the H3K27me3+H4K20me1 combination in J2s, which was associated with 803 genes.

**Table 2.** Distribution of histone modifications in relation to protein-coding genes.

| Histone modifications and combinations | Associated genes in eggs | Associated genes in J2s |
| --- | --- | --- |
| H3K4me3 | 6014 | 10564 |
| H3K9me3 | 1762 | 3475 |
| H4K20me1 | 1212 | 2127 |
| H3K27me3 | 829 | 1831 |
| H3K27ac | 756 | 1047 |
| H3K9me3+H4K20me1 | 531 | 699 |
| H3K27me3+H3K9me3+H4K20me1 | 362 | 571 |
| H3K27me3+H4K20me1 | 360 | 803 |
| H3K27ac+H4K20me1 | 250 | 299 |
| H3K4me3+H3K9me3 | 221 | 253 |
| H3K27ac+H3K27me3+H4K20me1 | 181 | 400 |
| H3K4me3+H4K20me1 | 150 | 221 |
| H3K27ac+H3K4me3 | 138 | 211 |
| H3K27me3+H3K9me3 | 96 | 220 |
| H3K27ac+H3K27me3 | 93 | 166 |
| H3K27ac+H3K27me3+H3K9me3+H4K20me1 | 66 | 92 |
| H3K27ac+H3K4me3+H4K20me1 | 57 | 105 |
| H3K4me3+H3K9me3+H4K20me1 | 42 | 35 |
| H3K27me3+H3K4me3 | 41 | 66 |
| H3K27ac+H3K9me3+H4K20me1 | 40 | 54 |
| H3K27ac+H3K9me3 | 26 | 79 |
| H3K27ac+H3K4me3+H3K9me3+H4K20me1 | 22 | 16 |
| H3K27ac+H3K4me3+H3K9me3 | 17 | 19 |
| H3K27ac+H3K27me3+H3K4me3+H4K20me1 | 14 | 28 |
| H3K27ac+H3K27me3+H3K4me3+H3K9me3+H4K20me1 | 13 | 22 |
| H3K27ac+H3K27me3+H3K9me3 | 8 | 31 |
| H3K27ac+H3K27me3+H3K4me3 | 7 | 10 |
| H3K27me3+H3K4me3+H4K20me1 | 5 | 8 |
| H3K27me3+H3K4me3+H3K9me3+H4K20me1 | 4 | 7 |
| H3K27me3+H3K4me3+H3K9me3 | 3 | 4 |
| H3K27ac+H3K27me3+H3K4me3+H3K9me3 | 2 | 7 |
| Total | 13322 | 23470 |

Numbers of *M. incognita*'s annotated protein-coding genes associated with H3K4me3, H3K9me3, H3K27ac, H3K27me3 and H4K20me1 histone modifications and their combinations, for Egg and J2 samples. Protein-coding genes were considered to be associated with a histone modification if at least 1 bp of the protein-coding gene annotation overlapped with the identified histone modification.

We then assessed the impact of each histone modification on gene expression (Fig 4). According to ChIP-seq and RNA-seq data, 10,242 genes in eggs and 18,577 genes in J2s were both expressed and associated with at least one histone modification.

**Fig 4.**
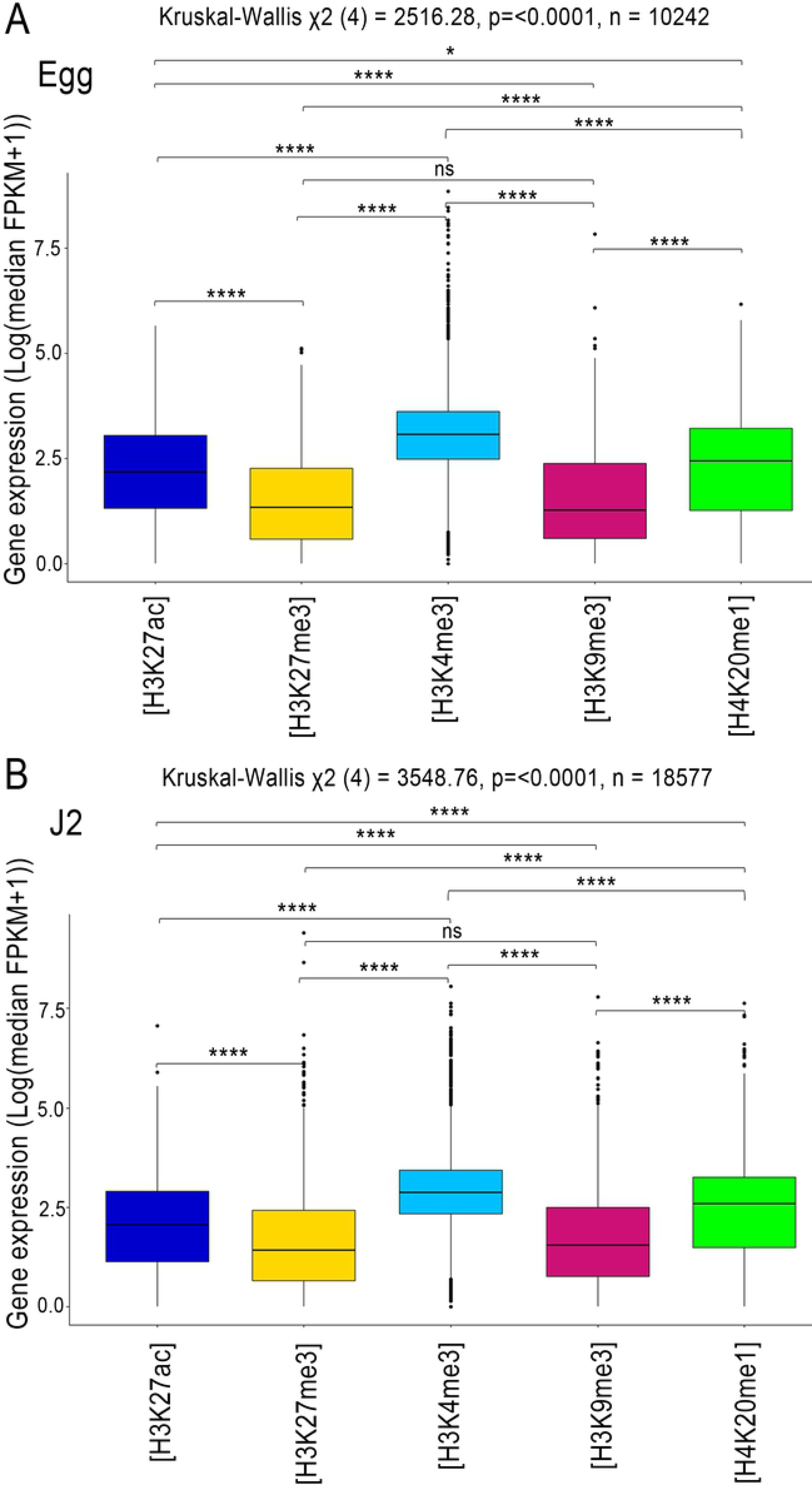
RNA-seq regulation of the protein-coding genes associated with histone modifications. Analysis of transcript levels of the genes associated with H3K27ac (blue), H3K27me3 (yellow), H3K4me3 (sky blue), H3K9me3 (dark pink) and H4K20me1 (green) in (A) Eggs and (B) J2s. Genes were considered to be associated with a histone modification if at least 1 bp of the annotation overlapped with an identified histone modification. Three biological replicates of *M. incognita* eggs and J2s have been treated jointly to identify common histone modification enrichment using ChromstaR. Normalized expression (Log(median FPKM+1) of genes, calculated triplicates is shown. A Kruskal-Wallis test was performed, followed by a pairwise Dunn test, to assess the significance of differences in gene expression level between the 5 histone modifications. ns *p*> 0.05, * *p*≤0.05, ** *p*≤0.01, \*\*\**p*≤0.001, **** *p*≤0.0001.

The distribution of gene expression values was shifted towards the highest median values for H3K4me3, and toward the lowest median values for the histone modifications H3K9me3 and H3K27me3 (Fig 4A-B). The other two known histone modifications, H3K27ac and H4K20me1 modifications, were associated with intermediate levels of gene expression (Fig 4A-B). These observations are consistent with observations for *C. elegans,* in which euchromatic regions with active transcription are enriched in H3K4me3 and H3K27ac, whereas regions with low levels of transcription activity are enriched in H3K9me3 and H3K27me3 [30].

We analyzed the top and bottom 10% of protein-coding genes ranked according to expression levels, to explore the proximal regulatory elements. We extended the area of overlap considered to 2 kb upstream and downstream from the protein-coding genes, with ChromstaR (Fig 5). For the top 10% of expressed genes at both stages, H3K4me3 enrichment overlapped start codon, implying that this histone modification occurs preferentially at the TSS of highly expressed genes (Fig 5A and Fig 5C). H3K27ac presented a “flat” profile, indicating a lack of evident enrichment. For H3K27me3, H3K9me3 and H4K20me1, the log(expected/observed) value was below zero, indicating that the most strongly expressed genes were depleted in these histone modifications. By contrast, the enrichment profile of H3K4me3 in the 10% of genes with the lowest levels of expression appeared as a “valley”, indicating depletion (Fig 5B and Fig 5D). The H3K27ac, H3K9me3, H3K27me3 and H4K20me1 signals were flat between and around the genes (Fig 5B and Fig5D).

**Fig 5.**
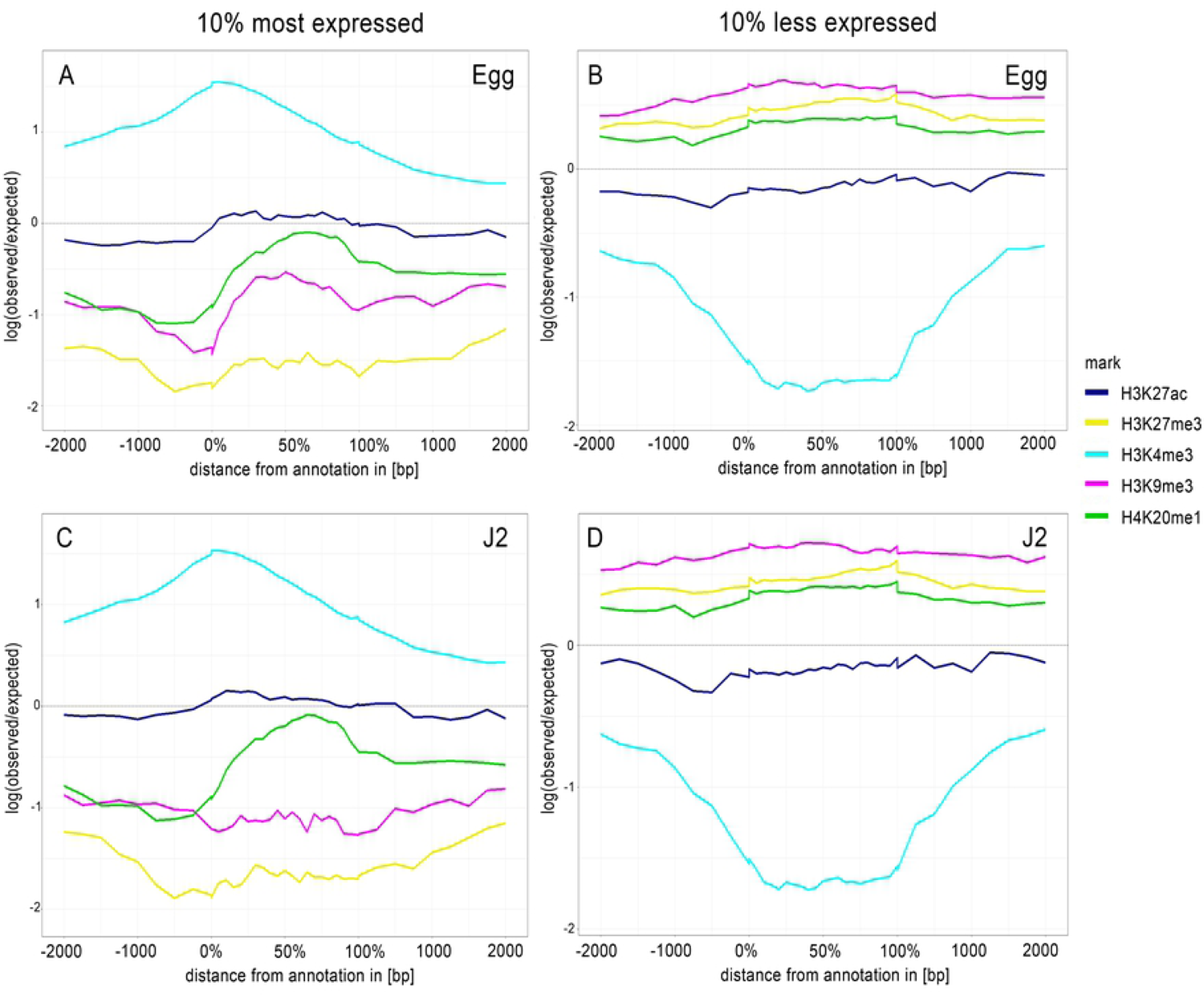
Average H3K4me3 enrichment profiles correlate with TSS regions of 10% most expressed genes. Average enrichment profiles of 5 histone modifications along a 4 kb region framing expressed protein-coding genes after ChIP-Seq of (A and B) eggs and (B and D) J2s. Average enrichment profiles were generated by ChromstaR, (log(observed/expected) value ranging from 1 (highly enriched) to −2 (depletion)), for 5 histone modifications: H3K27ac (blue), H3K27me3 (yellow), H3K4me3 (sky blue), H3K9me3 (dark pink) and H4K20me1 (green). Three biological replicates of *M. incognita* eggs and J2s have been treated jointly to identify common histone modification enrichment. Enrichment was analyzed for the (A and C) top and (B and D) bottom 10% of associated protein-coding genes ranked on the basis of RNA-seq normalized expression data (Log(median FPKM+1). *x-axis*: % in gene bodies and distance in bp upstream of TSS or downstream of TES. *y-axi*s: Density of mapped reads (log(observed/expected).

Finally, during the eggs-to-J2s transition, a change in the distribution of H3K4me3 was observed, with this modification disappearing from the TSS of underexpressed genes and becoming enriched at the TSS of overexpressed genes. This change in the distribution show a dynamic in histone modifications. However, it was less straightforward to establish a direct correlation between gene expression levels and the presence/absence of other histone modifications. The pattern of association between histone modifications and annotated protein-coding genes was, therefore, robust only for H3K4me3, and was associated with an expression switch during the eggs-to-J2s transition.

### Stage-specific enrichment in GO terms for genes associated with H3K4me3

Given the strong association of H3K4me3 with the higher expression of protein-coding gene expression, we compared functional annotations in eggs and J2s. We identified 6,014 genes in eggs and 10,564 genes in J2s as associated with H3K4me3. We then annotated the corresponding *M. incognita* proteins thanks to Interproscan [21]. Enrichment was detected for 46 GO terms in eggs and 8 GO terms in J2s (Fig 6). GO terms such as “ribonucleoside- and nucleoside-associated processes” were associated with the egg stage, whereas compounds identified in the metabolic processes’ ontology such as “ether”, “citrate” and “tricarboxylic acid” were specifically enriched in the J2 stage. We also identified 40 GO terms as displaying enrichment at both stages, with the strongest enrichment observed for processes related to protein biosynthesis: “translation”, “peptide biosynthetic process”, “peptide metabolic process”, “amide biosynthetic process”, “cellular amide metabolic process” (Fig 6).

**Fig 6.**
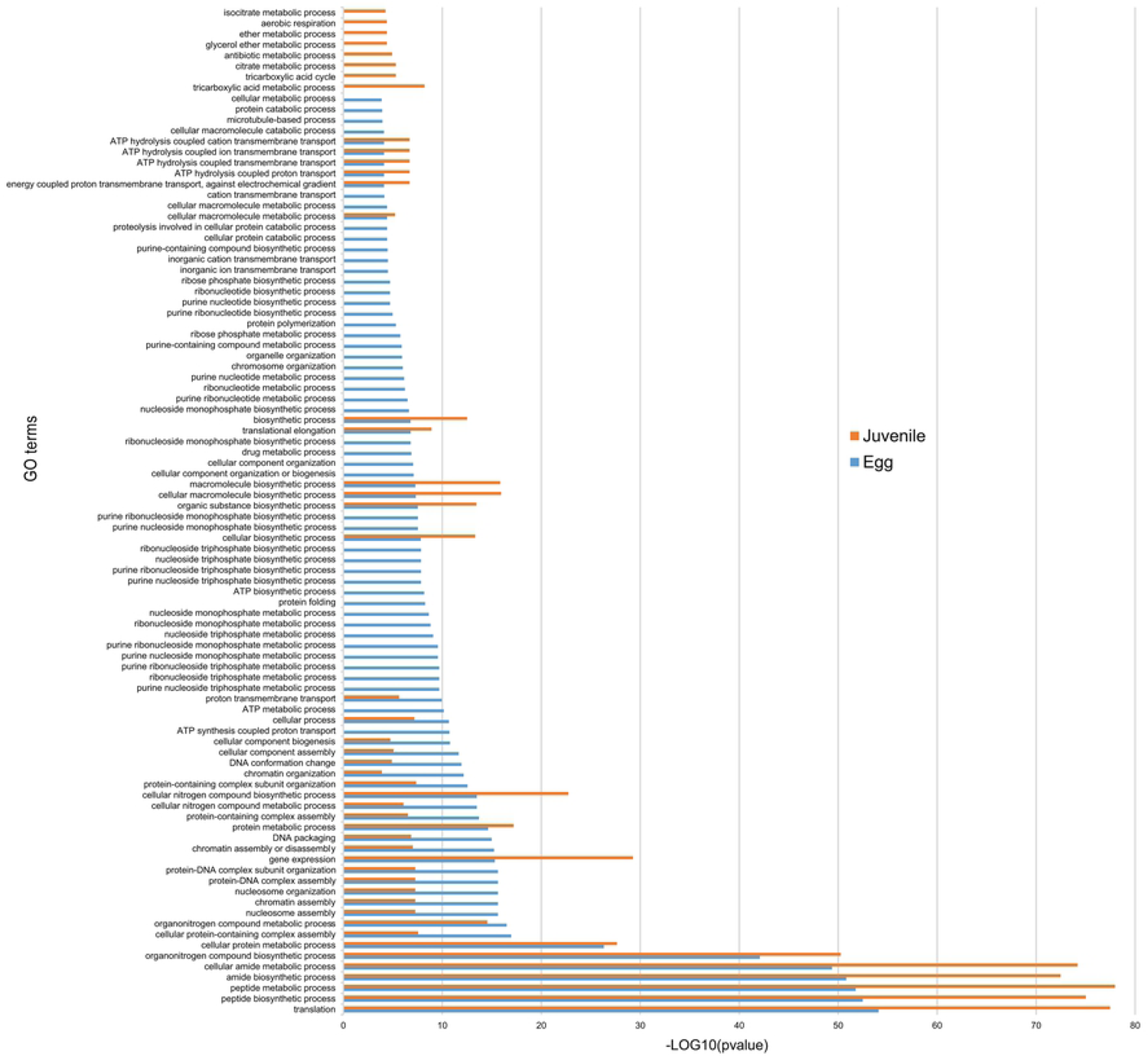
Functional annotation of protein-coding genes associated with H3K4me3. Histogram showing the ‘Biological processes’/Gene ontology (GO) term enrichment of protein-coding genes associated with H3K4me3. Protein-coding genes were considered to be associated with H3K4me3 if at least 1 bp of the protein-coding gene annotation overlapped with this histone modification. Three biological replicates of *M. incognita* eggs and J2s have been treated jointly to identify common histone modification enrichment. Overrepresented GO terms, in eggs (blue; 6,014 genes) and J2s (orange, 10,564 genes), were identified with GoFuncR with a FWER < 0.05 cutoff. X-axis is the -log10(pvalue) calculated to represent GO term enrichment on the bar chart.

Our observations of H3K4me3 dynamics during the eggs-to-J2s transition led us to analyze the functions of the products of the differentially expressed genes. We identified 89 genes in eggs and 177 genes in J2s as both associated with H3K4me3 and differentially expressed. Overrepresentation was detected for 39 GO terms specific to eggs (56/89 genes), and 9 GO terms specific to J2s (28/177 genes). GO terms linked to genomic organization and cell cycle-associated processes were associated with the egg stage, whereas cell signaling, and stimulus responses were specific to the J2 stage (Fig 7).

**Fig 7.**
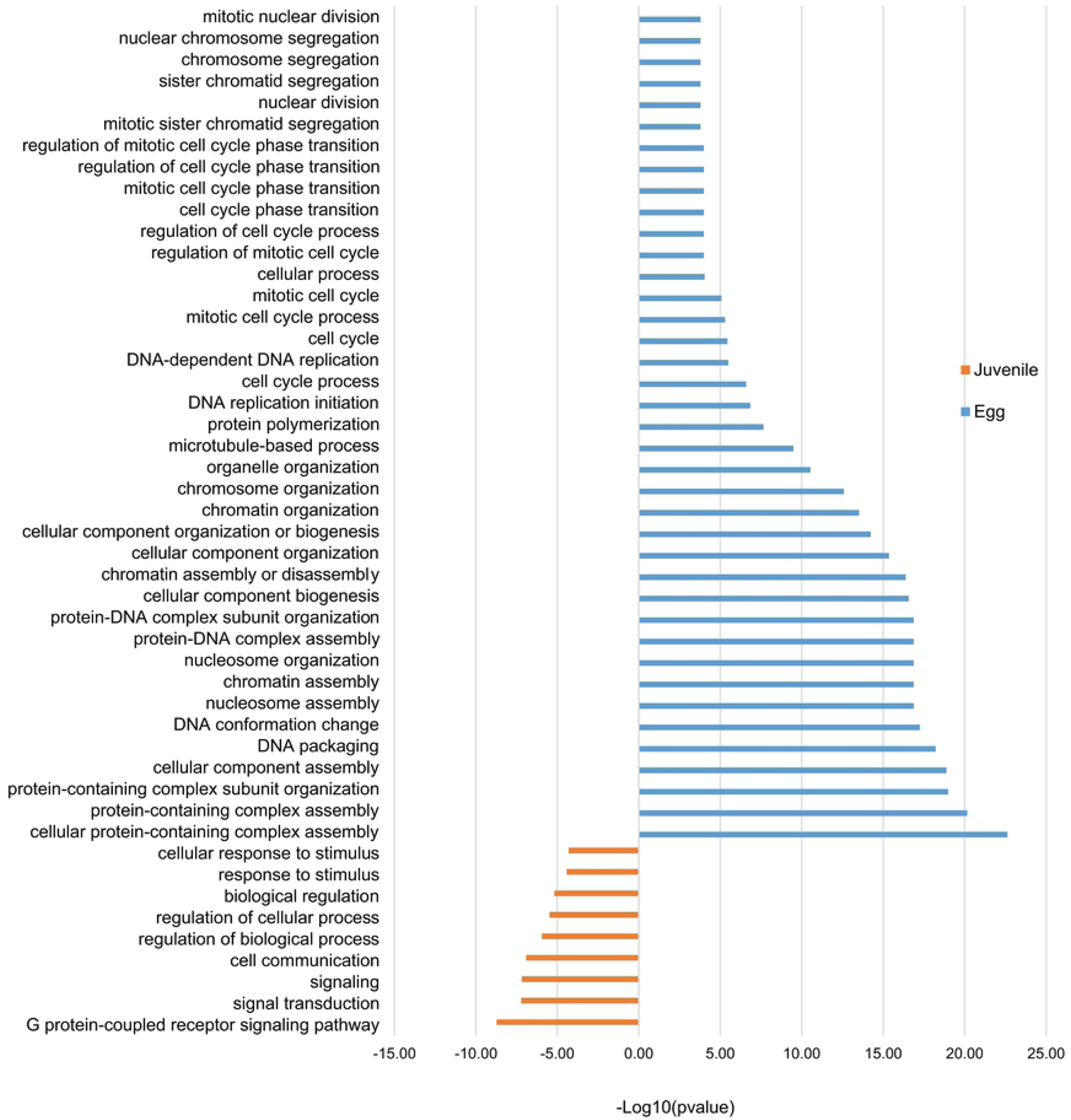
Stage-specific enrichment in Gene Ontology (GO) terms for protein-coding genes associated with H3K4me3. Histogram showing the “Biological processes’’/GO term enrichment of protein-coding genes showing both (i) differential gene expression during eggs-to-J2s transition; and (ii) H3K4me3 association. Differential gene expression was calculated using DESeq2 on triplicates, with a p value < 0.05 as a threshold for overexpression. Protein-coding genes were considered to be associated with H3K4me3 if at least 1 bp of the protein-coding gene annotation overlapped with this histone modification. Three biological replicates of *M. incognita* eggs and J2s have been treated jointly to identify common histone modification enrichment. Overrepresented GO terms, in eggs (blue; 89 genes) and J2s (orange, 117 genes), were identified with GoFuncR. with a FWER < 0.05 cutoff. X-axis is the -log10(pvalue) calculated to represent GO term enrichment on the bar chart.

The identification of orthologs in *C. elegans* and parasitic nematodes provided insight into the functions of the H3K4me3-associated genes differentially expressed during the eggs-to-J2s transition. We found 63 genes in *M. incognita* eggs and 119 genes in J2s for which at least one ortholog was present in *C. elegans*. Interestingly, orthologs of genes linked to the regulation of histones, DNA metabolism, cytoskeleton organization and the mitotic checkpoint were overrepresented among the most expressed genes in *M. incognita* eggs (Table S2). In *M. incognita* J2s, we identified orthologous genes involved in redox status and the regulation of cell trafficking (Table S3).

**Table 3.**
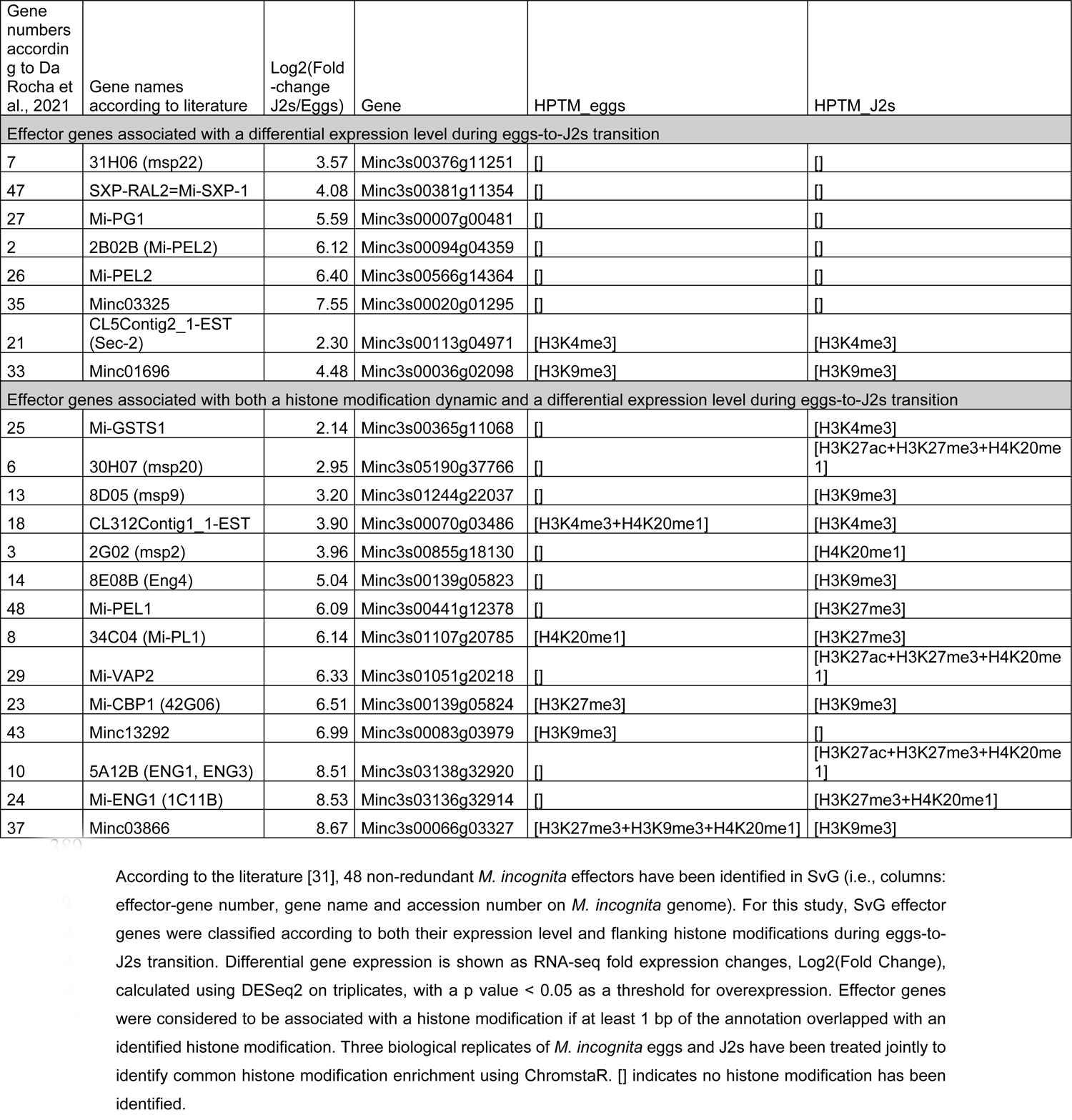
Transcriptional regulation of known subventral glands (SvG) effector genes.

### RKN effector-coding genes are subject to regulation by histone modifications

Effectors are secreted proteins that are essentials to nematode parasitism. Studies using RNA-Seq technology provided a comprehensive transcriptome profiling of *M. incognita* effector-producing glands and an opportunity to characterize their different patterns on infective aptitude, from the penetration to the successful interaction leading to feeding sites and the production of the next generation of eggs [31, 32]. In *M. incognita*, subventral glands (SvG) are mostly active during the earliest steps, whereas dorsal gland (DG) is active in the latest steps of the infection. A total of 48 and 34 putative non-redundant *M. incognita* effectors have been identified in SvG and DG, respectively [31]. We looked for histone modification associated with effector genes that are overexpressed in J2s. Among the 48 SvG effectors, 14 were associated with both a histone modification dynamic and a differential expression pattern during eggs-to-J2s transition. Only two of those effectors, Mi-GSTS1 and msp2, showed an appearance of activating histone modification in J2s. In contrast, combinations of histone modifications involving the repressive modifications H3K27me3 and H3K9me3 appeared to be the most abundant in this class of effectors (Table 3, S4 Table). Among the 34 DG effectors, 4 were associated with both a histone modification dynamic and a differential expression pattern during eggs-to-J2s transition. All of them were associated with combinations of repressive histone modifications (Table 4, S5 Table). Interestingly, the Mi-14-3-3-b DG effector exhibits reverse dynamics during the transition from eggs to J2s with a repression of expression in J2s associated with the appearance of H3K27me3. Altogether, these results suggest that histone modifications act as crucial regulators to precisely produce some effectors in a dose manner and in temporal sequence during parasitism.

**Table 4.** Transcriptional regulation of known dorsal gland (DG) effector genes.

| Gene numbers according to Da Rocha et al., 2021 | Gene names according to literature | Log2(Fold-change J2s/Eggs) | Gene | HPTM_egg | HPTM_juvenile |
| --- | --- | --- | --- | --- | --- |
| Effector genes associated with a differential expression level during eggs-to-J2s transition |  |  |  |  |  |
| 28 | Minc12639 | 3.18 | Minc3s00340g10545 | [] | [] |
| 25 | Minc01595 | 3.90 | Minc3s01184g21493 | [H3K27me3+H4K20me1] | [H3K27me3+H4K20me1] |
| Effector genes associated with both a histone modification dynamic and a differential expression level during eggs-to-J2s transition |  |  |  |  |  |
| 12 | 34F06 (msp24) | 2.68 | Minc3s00321g10151 | [H3K27me3] | [H3K27me3+H3K9me3] |
| 26 | Minc02097 (35A02, msp25) | 4.80 | Minc3s00202g07465 | [] | [H3K27me3] |
| 9 | 25B10 (msp33) | 4.99 | Minc3s03649g34419 | [] | [H3K9me3] |
| 16 | 6F06 (msp4) | 5.65 | Minc3s02324g29465 | [H3K9me3] | [] |
| 23 | Mi-14-3-3 | -3.43 | Minc3s00122g05244 | [] | [H3K27me3] |

## Discussion

Many biological processes involve chromatin changes, including the delimitation of functional elements in the genome and transcription regulation, particularly during complex parasitic life cycles. The RKN *M. incognita* has a wide host range and is found worldwide. Despite its clonal reproduction, *M. incognita* can rapidly adapt to unfavorable conditions [7, 8]. Epigenetic mechanisms may contribute to this rapid adaptation and the parasitic success of this nematode. Cytosine methylation is absent, or present at only very low levels, in the *M. incognita* genome, which contains no genes encoding DNA methyltransferases [20, 21]. Conversely, histone (de)acetylation and (de)methylation enzymes are present and conserved in the genome of *M. incognita* [21]. However, the role of histone modifications in phytoparasitic nematode biology remains unknown. We deciphered the chromatin landscape in the RKN *M. incognita* by studying five histone modifications and analyzing their dynamics during development. These modifications were not randomly distributed in *M. incognita* and colocalized with genomic elements, forming specific epigenetic signatures.

In the model nematode *C. elegans*, H3K4me3 enrichment is observed in actively expressed regions and therefore associated with euchromatin. By contrast, H3K9me3 and H3K27me3 enrichment is observed in silent genes, transposons, and other repetitive sequences and as such associated with heterochromatic regions. This histone code is observed in most organisms [33, 34]. However, these histone modifications do not have the same biological implications in some organisms [35]. H3K4me3 activates gene expression by a charge-mediated decompaction of the chromatin at promoter sequences [36]. H3K4me3 is usually distinguished by sharp peaks or enrichment around the TSS [37]. In *M. incognita*, we observed a typical profile of this type, with the sharp H3K4me3 peaks overlapping with the TSS of annotated protein-coding genes, associated with higher levels of gene expression. The conservation of the canonical function of H3K4me3 in *M. incognita* will pave the way for deciphering transcriptional regulation during development and parasitism.

Another activating histone modification, H4K20me1, typically results in diffuse chromatin modifications [38]. In *M. incognita*, H4K20me1 displayed a diffuse profile of this type over the entire genome and was associated with higher levels of expression than the repressive histone modifications. H4K20me1 levels have been shown to change dynamically during the cell cycle, peaking during the G2/M phase [39]. At the whole-organism scale, dynamic changes in H4K20 methylation have been observed during mouse preimplantation development, with this modification playing a key role in the maintenance of genome integrity [40]. *M. incognita* seems to have the same histone modification machinery as model organisms, and we can, therefore, predict analogous functions for H4K20me1 in cell cycle regulation and the maintenance of genome integrity in this nematode.

Another well-described histone modification is the heterochromatin-associated modification H3K9me3 [41], which plays an important role in regulating gene expression [42] and is characterized by distinct peaks at protein-coding genes [43]. H3K9me3 is also associated with TEs, which require controlled repression to prevent chaotic transposition in the genome. This modification has been described as the principal regulator of these elements in mouse embryonic stem cells [43] and in *C. elegans* [33]. In *M. incognita*, H3K9me3 presented sharp peaks in the bodies of genes with low levels of expression relative to other histone modifications. Moreover, the majority of H3K9me3 modifications were found on annotated TE, suggesting that H3K9me3 represses the mobile elements of the genome and indicating that its canonical function is also conserved in *M. incognita*.

Different sets of histone modifications can account for gene expression [44]. For instance, a balance between H3K27 trimethylation and H3K27 acetylation has been shown to regulate gene expression in a dynamic manner [45]. H3K27me3 is a broadly distributed repressive histone modification that downregulates gene expression, as demonstrated during development and cell differentiation [46, 47]. By contrast, H3K27ac is an activating histone modification that may be broadly distributed [48] or display narrow peaks [49]. In *M. incognita*, H3K27me3 and H3K27ac were broadly distributed throughout the genome despite their dual effects. H3K27ac is usually found on enhancers and can be used to distinguish between active and poised enhancers [50]. H3K27ac was associated with genes displaying higher levels of expression than those associated with repressive modifications in *M. incognita*. As in mammals, studies of H3K27ac enrichment could potentially be used to predict enhancers in *M. incognita* on the basis of local chromatin structure. Other sets of histone modifications may fine-tune regulation of the chromatin landscape, but their identification was limited by antibody availability and specificity in *M. incognita*.

Different combinations of histone modifications can be colocalized, acting together as activators, repressors or in a bivalent manner [51]. For instance, gene expression levels have been shown to be regulated by the ratio of H3K27Me3 to H3K4Me3 modifications, leading to a bivalent outcome: repression or activation [51]. In *M. incognita*, H3K27me3 enrichment was observed at the 5’UTR, potentially accounting for the low levels of expression for the associated genes. The colocalization of H3K27me3 and H3K4me3 at the 5’ UTR suggests bivalency for these two modifications in *M. incognita*. H3K27me3 enrichment was also found within tRNA-genes, suggesting a role for this modification in tRNA regulation, consistent with the presence of H3K27me3 near the RNA polymerase III binding sites used for the synthesis of tRNA in human embryonic stem cells [52].

Our findings indicate that histone modification is conserved *in M. incognita* and defines a reproducible and consistent landscape. We therefore further investigated the dynamics of histone modifications during development and parasitism, by considering the egg and juvenile stages. This developmental transition constitutes a major change in the nematode environment. J2s hatch and are released into the soil, in which they begin their life as mobile entities, moving towards the plant roots. The soil is a radically different environment from the eggs, and the newly hatched J2s must therefore adapt very rapidly to this new environment. At the scale of the genome, we found that the histone modification profile and gene expression level remained relatively stable during development. However, dynamical changes were highlighted during the eggs-to-J2s transition in *M. incognita,* in analyses at gene level. We identified pathways relating to the cell cycle as overrepresented in eggs, promoting *M. incognita* development. By contrast, J2s presented pathways linked to stimulus responses, reflecting the needs of J2s following their release into the soil after hatching, consistent with previous observations [31]. Furthermore, based on *C. elegans* orthology, egg stage-specific genes were involved in cell division, whereas J2s-specific genes were mainly involved in redox status regulation, reflecting the environment shift during the *M. incognita* life cycle. The identification of such candidate genes in *M. incognita* highlights the involvement of histone modifications in nematode development and could lead to the identification of new targets for pest control.

Histone modifications also contribute to the parasitic success of many animal or plant parasites. Parasites possess an arsenal of molecules known as effectors, which promote infection success. Fungal species, such as *Fusarium graminearum* or *Leptosphaeria maculans*, are the principal plant-parasitic organisms displaying chromatin-based control of concerted effector gene expression at specific times during infection [53, 54]. In *Zymoseptoria tritici*, the H3K27me3 distribution dictates effector gene expression during host colonization, preventing the expression of these genes when not required [55].

The association of J2s-overexpressed effector-coding genes with histone modifications suggests that epigenetic regulation contributes to *M. incognita* parasitism. However, by contrast to what we observed for stage-specific genes, the overexpression of effectors in J2s was not associated with H3K4me3, whatever the secretory gland, SvG and DG. For effectors, overexpression in J2s is mainly associated with combinations of repressive histone modifications. The overexpression of effector-coding genes needed at a specific time point, such as cell wall degrading enzymes during juvenile stage which help to the penetration of the nematode into the root system, may be under strong and complex regulation. In that respect, having effector-coding genes under repressive histone modifications could help the nematode to fine-tune their expression in a spatio-temporal way during plant infection. For instance, different histone modification dynamics may account for the coordinated, yet slightly different, expression of 2 pectate lyase genes, Mi-pel-1, and Mi-pel-2, during *M. incognita* infection of roots. Indeed, while these 2 genes were both overexpressed in early J2s stage (with similar fold-changes), only Mi-pel-1 gene was associated with repressive H3K27me3. Potential release of this repressive histone modification may provide an explanation for the expression of Mi-pel-1 only at late J2s stage, as previously reported [56].

Another example is the Mi-14-3-3-b DG effector gene which was the only one overexpressed in eggs and showing appearance of a H3K27me3 modification during the eggs-to-J2s transition. This result correlates with what is already known about the expression pattern of Mi-14-3-3-b during *M. incognita* infection with an early expression in eggs, a strong repression in J2s and a late expression in the female stage [57].

More generally, these results suggest that the fine-tuning of effector production during parasitism could be achieved through either another activating histone modification, still to be studied, or a different process such as transcription factors (TFs) activation. Consistent with this, a putative cis-regulatory element “Mel-DOG” has been identified in *M. incognita* DG effector promoters [31]. This might be the missing activator switch for the expression of DG effector genes at specific stages during the lifecycle of the nematode, even if the associated TFs are yet to be discovered. To achieve precise and accurate regulation of effector-genes, TFs and histones modifications may work in a cooperative way.

## Conclusion

We describe here the chromatin landscape of a parasitic nematode, revealing a dynamic process during the life cycle. This pioneering study shows that *M. incognita* presents a histone modification similar to that of the model nematode *C. elegans*. Beyond model organisms, the epigenome arguably plays an important role in development and the regulation of parasitism. The next step will be to decipher the epigenetic response of *M. incognita* to environmental changes, such as host adaptation, in greater detail.

## Materials and Methods

### Biological materials

One-month-old tomato plants, *Solanum lycopersicum* (St Pierre), were inoculated with soil infested with *M. incognita*. Eggs were collected from tomato roots seven-week-old after infection, by grinding, sterilizing, and filtering, as previously described [58]. Extracted eggs were purified by centrifugation on a 30% sucrose gradient, washed and either stored at −80°C for subsequent experiments or kept in autoclaved tap water, at room temperature, for seven days, to produce juveniles J2s. Hatched J2s were collected by filtration and centrifugation (13,000 x *g*, 1 min) and stored at −80°C.

### Antibody screening

Commercially available antibodies raised against histones with posttranslational modifications were selected on the basis of two criteria: ChIP-grade and preferentially used in the model nematode *C. elegans* (Table S1). We assessed the specificity of each antibody in *M. incognita* by a two-step method combining western blotting and ChIP-titration, as described by Cosseau [26].

### Western blot

Nematodes were resuspended in a homemade extraction buffer (3% SDS, 10% sucrose, 0.2 M DTT, 1.25 mM sodium butyrate, and 62.5 M Tris/Cl pH 6.8) and crushed with a glass Dounce homogenizer for 2 minutes. Samples were sonicated (Vibra Cell™) three times, at 70% intensity, for 15 s each, with cooling on ice during the intervals. They were then boiled for 5 minutes at 99°C after the addition of Laemmli buffer (Cat. #1410737, Biorad). Proteins were separated by SDS-PAGE and transferred to a membrane with a Trans Blot TURBO (Biorad). The membrane was incubated for 1 h at 37°C in a homemade blocking buffer (50 mM NaCl, 0.05% Tween 20, 5% fat-free dry milk, 20 mM Tris/Cl pH 7) and then for 1.5 hours with antibodies in the blocking buffer. Finally, the membrane was washed and incubated with a secondary antibody. Signals were detected by incubation with SuperSignal West Pico Chemiluminescent Substrate (Thermo Fisher Scientific, Cat.34579) and to the use of ChemiDoc Imaging systems (Biorad). Antibodies that did not bind to a unique target were discarded from the analysis (S1 Table and S1 Fig).

### Crosslinking and chromatin immunoprecipitation (ChIP)

Frozen eggs or juveniles were resuspended in 500 µL Hank’s balanced salt solution (HBSS, Sigma-Aldrich, Cat. #H4641, Lot RNBG1861) and crushed with a glass Dounce homogenizer for 7 min. We then added 500 µL 1 x HBSS and transferred the solution to an Eppendorf tube. Samples were centrifuged (at 2,700 x *g*, 5 min, 4°C). For crosslinking, the pellet was resuspended in 1 mL 1 x HBSS containing 13.5 µL of 37% formaldehyde (Sigma-Aldrich, Cat. #252549), and incubated for 10 min at room temperature, with occasional inversion. Binding was stopped by adding 57 µL 2 M glycine (Diagenode, cat. C01011000) and incubating the sample for 5 min at room temperature. Samples were centrifuged at 2,700 x *g*, 4°C for 5 min. The pellet was rinsed twice, with 1 mL 1 x HBSS each, and centrifuged again (2,700 x *g*, 4°C for 5 min). ChIP was performed with the Auto-Chipmentation Kit for histones (Diagenode, cat. C01011000). Crosslinked chromatin was resuspended in 100 µL cold lysis buffer IL1 and incubated at 4°C for 10 min in a rotating well. Following centrifugation (2,700 x *g*, 4°C for 5 min), the supernatant was discarded, and the pellet was resuspended in 100 µL cold lysis buffer IL2, and incubated in a rotating well for 10 min at 4°C. The suspension was centrifuged (2,700 x *g*, 4°C for 5 min) and the pellet was resuspended in 100 µL of complete shearing buffer iS1. Samples were sonicated with the Bioruptor Pico, over 5 cycles (30 s ON and 30 s OFF). They were then transferred to new tubes and centrifuged (16,000 x *g*, 10 min, 4°C). The supernatants were transferred to new tubes, pooled by batch and 500 μL iS1 was added. The ChIPmentation program was selected on the Diagenode SX-8G IP-Star Compact. We used the following parameters: 3 hours of antibody coating at 4°C, 13 hours of IP reaction at 4°C, 10 min wash at 4°C and 5 min tagmentation. All steps were performed with the intermediate mixing speed.

ChIP-buffer, antibody coating mix and immunoprecipitation mix were prepared in accordance with the supplied protocol. Stripping, end repair and reverse cross-linking were performed with the reagents provided with the kit.

### Titration by qPCR

The immunoprecipitated DNA was quantified by qPCR with a LightCycler480 (Roche System). The PCR mix was prepared with 2 µL of immunoprecipitated chromatin, in a final volume of 10 µL (0.5 µL of each primer, 5 µL of Eurogentec Takyon™ SYBR® 2 x qPCR Mastermix Blue). The following Light-Cycler run protocol was used: denaturation at 95°C for 3 min; amplification and quantification (40 cycles), 95°C for 30s, 60°C for 30s, 72°C for 30s. Cycle threshold (Ct) was determined with the fit point method of LightCycler480 version 1.5. PCR was performed in triplicate, and the mean Ct was calculated.

Percent input recovery (%IR) was calculated as described by Cosseau [26], with the following formula: %*IR* = 100(*E^Ct(input)—Ct(IPbound)^*) where E is primer efficiency, Ct(input) is the Ct of the unbound fraction, and Ct(IPbound) is the Ct of the immunoprecipitated sample. Only antibodies reaching saturation were considered specific and were used for ChIP-Seq experiments, at their optimal concentration (S1 Table and S1 Fig).

### ChIP-Seq

The same ChIP protocol was performed with the Auto-Chipmentation kit for histones (Diagenode,cat. C01011000), with specific antibodies validated for *M. incognita,* targeting the histone modifications H3K4me3 (Merck Millipore ref 04-745, batch 2452485), H3K9me3 (Abcam ref ab8898, batch GR306402-2), H3K27ac (Abcam ref ab4729, batch GR150367-2), H3K27me3 (Epigentek ref A-4039, batch 503019) and H4K20me1 (Abcam ref ab9051, batch GR158874-1). For each antibody, ChIP was performed in biological triplicate on two different *M. incognita* stages: eggs and J2s. The ChIP control was the input-control, consisting of a fraction of sheared chromatin without immunoprecipitation [59].

Illumina libraries were constructed with primer indices provided by the Auto-Chipmentation kit for histones (Diagenode,cat. C01011000), according to the protocols supplied. The amount of DNA was determined and adjusted by qPCR quantification. Amplified libraries were quantified on a Bioanalyzer and sequenced by the BioEnvironnement platform (University of Perpignan, France) with an Illumina NextSeq 550 instrument generating 75 bp single-end reads. Sequencing reads have been deposited in the NCBI Sequence Read Archive (SRA, NCBI), under accession number PRJNA725801.

### ChIP-Seq data analysis

Graphical representations were generated, and statistical analyses were performed with R version 3.6.1 (www.r-project.org) and the following libraries: chromstaR, cowplot, bamsignals, gplots, reshape2, tidyverse, ggpubr and rstatix.

Illumina read quality was analyzed with FastQC [60]. Read trimming was performed with Trim Galore (http://www.bioinformatics.babraham.ac.uk/projects/trim_galore/), using the default parameters. Processed reads were mapped onto the reference genome of *M. incognita* [27] with Bowtie2, using “Very sensitive end-to-end” presets [61]. All library sizes were downsampled to the size of the smallest library we had, corresponding to 3.7 million reads.

Peak calling for domain visualization in the *M. incognita* genome was performed with Peakranger [62]. A fraction of sheared chromatin without immunoprecipitation has been used as input to subtract the background level. Normalized tracks were visualized with the Integrative Genome Viewer [63]. Biological replicates were treated independently, and reproducibility was checked manually (S2 Fig).

Enriched domain identification and chromatin state analysis were performed with ChromstaR, using the differential mode with default parameters, except for bin size and step size, which were set at 500 bp and 250 bp, respectively [28]. ChromstaR uses a hidden Markov model approach to predict domains displaying enrichment. The three biological replicates were treated jointly by ChromstaR to generate the HMM model. Peak prediction for each histone modification was defined by a posterior probability > 0.5.

The genomic frequencies of the histone modifications were calculated with ChromstaR and correspond to the percentage of bin sizes (∼500 bp) with histone modifications and their combinations (defined as the overlapping of multiple modifications on the same bin) over the 184 Mb of the *M. incognita* genome. As an example, H3K4me3 frequency (%) corresponds to the total covered bases (∼25MB) divided by the genome size (∼184MB) and multiplied by 100.

Analyses of enrichment at genomic elements were performed by plotting ChromstaR heatmaps. Heatmaps were generated from the logarithm(observed/expected) ratio. The “expected” parameter corresponds to the probability of a bin to be both a genomic element and marked with histone modification at the same location. The “observed” parameter constitutes the frequency of a bin corresponding to be both a genomic element and marked with histone modification at the same location. When the ratio is > 0, the genomic element is observed mor frequently than expected and considered as statistically enriched with the histone modification. We used genome annotation data from a previous genome sequencing analysis of *M. incognita,* including 43,718 protein-coding genes (corresponding to mRNA annotation) [27]. Furthermore, canonical TEs were annotated and filtered using REPET [10, 60]. Regions of differential enrichment were determined with a minimum differential posterior, to detect pairwise differences at *p*=0.9999.

### Transcription analysis and histone modification profile

RNA-seq data were provided by previous analyses of different life stages, eggs and J2s, of the nematode [27]. Data was reprocessed by Kozlowski and coworkers [10], to generate FPKM values. Raw FPKM values were transformed to obtain Log(median FPKM+1) values, keeping the median of the three biological replicates as a representative value. Raw FPKM values are available online [64]. The number of genes associated with histone modifications was calculated by determining whether the gene position overlapped a position of histone modification enrichment by at least 1 bp. A boxplot representing the levels of gene expression associated with the five histone modifications was generated for genes for which expression data were available. A Kruskal-Wallis test was performed, followed by a pairwise Dunn test, to identify significant differences in gene expression level between different histone modifications, with a *p* value < 0.05 was considered significant.

The mean enrichment profiles were calculated by ChromstaR, based on the log(expected/observed) enrichment from 2 kb upstream to 2 kb downstream from the protein-coding genes, considering only the top and bottom 10% of genes ranked according to expression level associated with the five histone modifications.

Differentially expressed genes were identified using previous RNA-seq data [27, 65] processed by DE-seq2 [66], a *p* value < 0.05 was considered significant and a fold-change > 2 for overexpression.

### GO enrichment analysis

GO term enrichment was analyzed with the R package GOfuncR, using default parameters. The FWER cutoff was set at 0.05 to identify overrepresented GO terms. (-)Log_10_(pvalue) was calculated for the representation of GO terms, more specifically “Biological Processes”. All *M. incognita* genes associated with GO terms were used as references for GO enrichment analysis (i) for genes associated with H3K4me3 only; and (ii) for genes both associated with H3K4me3 only and differentially expressed during eggs-to-J2s transition.

*M. incognita* orthologs were identified from a previous work [21] using FamilyCompanion to identify orthologous links with 20 other species. Searches for GO terms for *C. elegans* orthologs were performed with the functional classification included in the PANTHER webtool [67].

## Acknowledgments

We would like to thank Jean-François Allienne and the IHPE team (Perpignan, France) for ChIP sample preparation and sequencing. We thank Dr. Marc Bailly-Bechet, Dr. Dominique Collinet, Julie Dazenière, Dr. Georgios Koutsovoulos and Dr. Djampa Kozlowski (ISA Sophia Antipolis, France) for assistance with and discussions about data analysis and statistics. We also thank Yongpan Chen, Dr. Joffrey Mejias, Yara Nourredine, Laura Perrot and Salomé Soulé (ISA Sophia Antipolis, France) for their help in the preparation of biological materials and fruitful discussion. We also thank Dr. Michael Quentin (ISA Sophia Antipolis, France) for help with effector analysis. We thank all the members of the IPN team for insightful discussions and technical help. We also thank Corinne Rancurel and PlantBios’s BIG bioinformatics platform for technical support (ISA Sophia Antipolis, France).

## Supporting information

**S1 Figure. Two-step antibody validation for ChIP-Seq.** Examples of two-step antibody validation adapted from (Cosseau et al., 2009): Western blot detection of **(A)** acetylated H3 at lysine 27 (H3K27ac), **(B)** monomethylated H4 at lysine 20 (H4K20me1) and **(C)** trimethylated H3 at lysine 27 (H3K27me3). qPCR validation on immunoprecipitated chromatin from *M. incognita*, with various volumes (0-16 µL) of **(D)** anti-H3K27 acetyl and **(E)** anti-H4K20 monomethyl antibodies. The percent input recovery (%IR) was calculated from the targeted amount of DNA and normalized with respect to the percent input recovery for the housekeeping gene. **(A)** and **(D)** show examples of successful validation for both western blotting and ChIP-titration, whereas **(B)** and **(E)** were validated only on western blotting, and **(C)** was not validated at the first step.

**S2 Figure. H3K4me3, H3K9me3, H3K27ac, H27me3 and H4K20me1 histone modifications on the *M. incognita* genome.** Triplicate tracks of histone modifications are illustrated on *M. incognita* at high resolution. Sequence reads for (A) H3K4me3 (blue, scaffold Minc3s00004), (B) H3K9me3 (red, scaffold Minc3s00013), (C) H3K27ac (pink, scaffold Minc3s00038), (D) H3K27me3 (green, scaffold Minc3s00003) and (E) H4K20me1 (black, scaffold Minc3s00007) samples were visualized in IGV software. Values shown on the y axis represent the relative enrichment of ChIP-Seq signals obtained with PeakRanger (peaks correspond to read counts after background/input subtraction). For each histone modification, the three biological replicates (rep1, rep2 and rep3) are shown. Each track contains information from one biological replicate of eggs.

**S3 Figure. Genome wide distribution of histone modifications, J2 samples, in relation to annotations for the *M. incognita* genome.** The distribution of histone modifications was analyzed with ChromstaR, which calculated the spatial enrichment in histone modifications for different available genomic annotations. Enrichment is represented as the log(observed/expected) value and ranges from 2 (highly enriched, red) to −2 (poorly enriched, blue). This enrichment heatmap is a 5×10 matrix representing 5 histone modifications (H3K4me3, H3K9me3, H3K27ac, H3K27me3 and H4K20me1) and 10 genomic annotated elements (CDS, exon, five prime UTR, gene, mRNA, ncRNA, rRNA, TE, three prime UTR and tRNA). Three biological replicates of *M. incognita* J2s have been treated jointly to identify common histone modification enrichment.

**S4 Figure. Distinct epigenetic landscapes at the transposable element (TE) and protein coding-gene annotations in *M. incognita*.** (A) Illustration of H3K9me3 enrichment in association with TE. Screenshot of the full scaffold Minc3s00875 with selected tracks for H3K9me3 (red), TE and gene annotations (dark blue). (B) Illustration of H3K4me3 enrichment in association with expressed protein-coding genes. Screenshot of the full scaffold Minc3s03894 with selected tracks for H3K4me3 (sky-blue), gene annotations (dark blue) and eggs transcripts (RNA-seq; grey). Samples were visualized in IGV software. Values shown on the y axis represent the relative enrichment of ChIP-Seq signals obtained with PeakRanger (peaks corresponding to read counts, normalized by the percent input method). Each track contains information from one biological replicate of eggs.

**S1 Table. Antibodies selected and tested for ChIP-seq analysis on *M. incognita*.** Antibodies were selected based on their availability, their validation in *C. elegans* and their validation as ChIP-grade when possible. Each antibody was tested by a two-step validation process, as described by Cosseau et al., 2009. The amount of antibody used was determined by titration at the saturation point.

**S2 Table. *M. incognita* Egg-overexpressed genes orthologs in *C. elegans*.** GO enrichment analysis showed specific terms associated with orthologous egg-overexpressed genes. Protein family/subfamily and classes were obtained using PANTHER and detailed for each *M. incognita* and *C. elegans* gene, with their GO ID.

**S3 Table. *M. incognita* J2-overexpressed genes orthologs in *C. elegans*.** GO enrichment analysis showed specific terms associated with orthologous J2-overexpressed genes. Protein family/subfamily and classes were obtained using PANTHER and detailed for each *M. incognita* and *C. elegans* gene, with their GO ID.

**S4 Table Transcriptional regulation of known subventral glands (SvG) effector genes.** According to the literature [31], 48 non-redundant *M. incognita* effectors have been identified in SvG (i.e., columns: effector-gene number, gene name and accession number on *M. incognita* genome). For this study, SvG effector genes were classified according to both their expression level and flanking histone modifications during eggs-to-J2s transition. Differential gene expression is shown as RNA-seq fold expression changes, Log2(Fold Change), calculated using DESeq2 on triplicates, with a p value < 0.05 as a threshold for overexpression. Effector genes were considered to be associated with a histone modification if at least 1 bp of the annotation overlapped with an identified histone modification. Three biological replicates of *M. incognita* eggs and J2s have been treated jointly to identify common histone modification enrichment using ChromstaR. [] indicates no histone modification has been identified. NS indicates no difference in gene expression between egg and J2 samples. NA indicates no predicted genes on *M. incognita* genome.

**S5 Table. Transcriptional regulation of known dorsal gland (DG) effector genes.** According to the literature [31], 34 non-redundant *M. incognita* effectors have been identified in DG (i.e. columns: effector-gene number, gene name and accession number on *M. incognita* genome). For this study, DG effector genes were classified according to both their expression level and flanking histone modifications during eggs-to-J2s transition. Differential gene expression is shown as RNA-seq fold expression changes, Log2(Fold Change), calculated using DESeq2 on triplicates, with a p value < 0.05 as a threshold for overexpression. Effector genes were considered to be associated with a histone modification if at least 1 bp of the annotation overlapped with an identified histone modification. Three biological replicates of *M. incognita* eggs and J2s have been treated jointly to identify common histone modification enrichment using ChromstaR. [] indicates no histone modification has been identified. NS indicates no difference in gene expression between egg and J2 samples. NA indicates no predicted genes on *M. incognita* genome.

## Notes

**Funding:** This work was supported by INRAE and the French Government (National Research Agency, ANR) through the ANR-18-CE20-0002 (ADMIRE), LABEX SIGNALIFE ANR-11-LABX-0028 and IDEX UCAJedi ANR-15-IDEX-0 programs. With the support of LabEx CeMEB, an ANR « Investissements d’avenir » program (ANR-10-LABX-04-01). This study is set within the framework of the « Laboratoire d’Excellence (LabEx) » TULIP (ANR-10-LABX-41).

### Competing Interest Statement

The authors have declared no competing interest.

